# Genetic association between angiotensin converting enzyme gene in Saudi obese women established gestational diabetes; Non-replication of case-control studies in insertion and deletion polymorphism

**DOI:** 10.1101/499137

**Authors:** Abdulaziz A. Alodhayani

## Abstract

**Introduction:** Gestational diabetes mellitus (GDM) is defined as carbohydrate intolerance detected during pregnancy and resolves after delivery. Presently, GDM was identified with biochemical but not with molecular analysis. Therefore, this study investigated the genetic, biochemical, clinical association between *ACE* polymorphism in GDM women. Methods: In this case-control study, 400 pregnant women were selected to inspect the *ACE* genotyping. Two hundred GDM cases were compared with 200 non-GDM subjects. Genotyping was carried out with direct primer sequence followed with 3% agarose gel electrophoresis.

**Results:** Statistical association was identified between GDM and non-GDM subjects with D allele, ID genotype and dominant model (D vs I: OR-1.62 [95%CI: 1.09–2.41]; p=0.01; ID vs II: OR-1.58 [95%CI: 1.02–2.42]; p=0.03; DD+ID vs II; OR-1.62 [95%CI: 1.09–2.40]; p=0.01). when comparison was carried out in obese and non-obese of GDM subjects' negative association was confirmed (p>0.05) and Weight cum BMI associated positively with *ACE* II, ID and DD genotypes (p=0.07 and p=0.02).

**Conclusion:** This results confirm positive association with *ACE* variant in GDM subjects; confirms the association between GDM and obesity in Saudi women. However, future studies should be implement in other diseases in Saudi as well as GDM with obesity studies in global populations to rule out the diseases.

## Introduction

Pregnant condition of women with undiagnosed diabetes reveals varying degree of glucose intolerance is defined as gestational diabetes mellitus (GDM), globally effecting 1.1%-14.3% during pregnancy and has ~35.6%-69% of recurrent risk.^1^ Earlier studies have addressed ~3–65% of GDM women developed type 2 diabetes mellitus (T2DM) during 5–16 years after the index pregnancy.^2^ Human and Animal studies research revealed GDM and T2DM have characterized β-cell dysfunction, insulin resistance and defects in insulin action.^3^ T2DM and GDM share the common pathophysiology and predisposing factors.^4^ Family history of GDM is one of the major risk factor for the pregnant women to prone chronic diseases as T2DM, obesity (weight gain) and hypertension.^5^ Adapting the western life style with uncontrolled diet, may be obese during childhood and epigenetic changes could be the main cause for weight gain.^6^ A significant risk factor for GDM has been documented as maternal health, macrosomia and cesarean delivery.^7^ Born neonates to GDM mother will have risk of low blood glucose other complications such as jaundice, pre-eclampsia, polycythemia and respiratory related syndromes.^8^ All types of diabetes such as T1DM, T2DM, GDM and specific types of diabetes due to other causes has genetic com--ponent. However genetic knowledge in diabetes accumulating progressively in diabets etiology and potential treatment of our updated knowledge in diabetes was not utilizing opportunities for modified management of diabetes in pregnancy.^9^ Importance of single nucleotide polymorphisms (SNPs) studies in coding and non-coding regions of specific genes was to understand the pathophysiology of human diseases for improving diagnosis, prevention, treatment and disease-causing genes. Approximately, 90% of sequence variants in humans are in DNA sequences that code for proteins.^10^ Genetic scientists have documented the significant progress in detecting genetic diseases.^11^ Angiotensin converting enzyme (ACE) gene is one of the important biomarker identifying in all multifactorial diseases mainly diabetes as T1DM, T2DM, GDM, New onset of diabetes after transplantation, diabetic nephropathy and retinopathy.

Previous studies have exposed as Renin angiotensin system may play an important role in T2DM nephropathy.^12^ *ACE* gene functional polymorphism consist of Insertion and deletion (I/D) was identified with ACE plasma levels, first genetic variant connected with human physical performance are strong linkage disequilibrium with genetic factors.^13^ It appears on chromosome 17q23 with 26 exons, 25 introns which are 44,770 bp in size and on 16^th^ intron I allele consist of 287 bp of non-coding Alu repeat sequence is absent in D allele, confirms II, ID and DD genotypes.^14, 15^ However, limited studies have been implemented in GDM disease as well as in Saudi population. Therefore, this study aims to investigate the insertion and deletion polymorphism in the ACE gene in Saudi women effected with GDM.

## Materials and methods

### Saudi subjects

Ethical approval was the initial step which was granted from King Saud University (KSU). This case-control study was designed from Department of obstetrics and gynecology. Altogether, 400 Saudi subjects signed the consent form were recruited based on inclusion and exclusion criteria.^16^ and divided as GDM (*n*=200) and non-GDM (*n*=200), also termed as control subjects. Overall, 5 ml of the peripheral white blood cells were collected from each patient and divided them into (a) coagulant blood (3 ml) was used for the reconfirmation of GDM as well as for lipid analysis and (b) 2 ml of the EDTA blood was used for the separation of genomic DNA for molecular analysis.^17^

### Anthropometric and Biochemical analysis

Saudi women’s Body Mass Index (BMI) were calculated with recorded height and weight by well trained-experienced female nurses. Age, mean gestational age, Hypertension and family history was recorded. Screening of glucose intolerance was performed in all the pregnant women (*n*=400) between 24–28 gestational weeks. Glucose challenge test (GCT) was initial preliminary screening test carried out and abnormal GCT was processed for oral glucose tolerance test (OGTT) with 100 grams of glucose. Positive turned subjects were considering as GDM and negative subjects were as non-GDM. Glucose analysis confirmation was carried out with American Diabetes Association Criteria. The details of GCT and OGTT values and protocol were described in Al-Hakeem et al^17^ study. Remaining serum sample was used for lipid profile analysis. Separately, fasting and post-prandial blood parameters were measured for biochemical analysis.

### Molecular analysis

AccuVis Bio DNA kits were used to extract the genomic DNA from white cells and extracted DNA was quantified with NanoDrop to check the quality of DNA. The forward and reverse oligonucleotides and PCR amplification conditions were selected from earlier study^14^,direct PCR was implemented for genotyping and successful amplification was confirmed with 3% agarose gel. Insertion band indicates 490bp and deletion band indicates 190bp; However, heterozygous band indicates 490 and 190 bp respectively.

### Statistical analysis

Anthropometric, biochemical and clinical characteristics between GDM and non-GDM subjects were expressed as mean±standard deviation. Genotype, allele frequencies and chi-square (χ^2^), 95% confidence intervals (CI) and p values were calculated within the Saudi subjects. Gene counting protocol was followed for calculating the alleles and Hardy-Weinberg equilibrium (HWE) was resoluted to test the *ACE* alleles. Anova analysis for variance was also studied for genotype and clinical details.

## Results

### Non-genetic analysis

Baseline characteristics of GDM and non-GDM subjects of anthropometric, biochemical and clinical details are tabulated in Table 1. GDM subjects were elder than control subjects comparing with the age, height, weight and BMI. Significant difference was observed with age, weight and BMI (p<0.05). As shown in Table 1, the biochemical parameters as FBS, PPBG, GCT, OGTT (fasting, 1^st^, 2^nd^ and 3^rd^ hours) and lipid profile such as TC, TG, HDL-C (but not LDL-C) were statistically associated with GDM comparing to non-GDM subjects (p<0.05). Family histories of T2DM and GDM were shown to be statistical significant (p<0.05).

**Table 1:** Clinical features of pregnant women ripens with and without diabetes

| Characteristics | GDM Cases<br>(n=200) | non-GDM<br>(n=200) | p value |
| --- | --- | --- | --- |
| Age (Years) | 32.4±5.7 | 31.3±6.0 | 0.04 |
| Weight (Kg) | 77.1±13.3 | 74.8±12.0 | 0.007 |
| Height (cm) | 158.5±5.9 | 157.8±5.3 | 0.16 |
| BMI (kg/m<sup>2</sup>) | 34.4±4.6 | 33.3±4.2 | 0.006 |
| Gestational Age (weeks) | 30.27±5.7 | NA | NA |
| FBS (mmol/L) | 5.0±0.9 | 4.5±0.8 | 0.0001 |
| PPBG (mmol/L) | 6.8±2.0 | 4.9±1.8 | 0.0001 |
| GCT (mmol/L) | 9.5±1.8 | 6.3±1.5 | 0.0001 |
| OGTT (Baseline) (mmol/L) | 5.2±1.1 | 4.5±0.8 | 0.0001 |
| OGTT (1<sup>st</sup> hour) (mmol/L) | 10.7±1.8 | 8.0±1.7 | 0.0001 |
| OGTT (2<sup>nd</sup> hour) (mmol/L) | 9.2±1.8 | 6.7±1.6 | 0.0001 |
| OGTT (3<sup>rd</sup> hour) (mmol/L) | 5.6±1.7 | 4.5±1.3 | 0.0001 |
| TG (mmol/L) | 2.3±1.8 | 1.7±0.9 | 0.0001 |
| TC (mmol/L) | 5.7±1.2 | 5.2±1.0 | 0.0001 |
| HDL-C (mmol/L) | 0.91±0.3 | 0.64±0.2 | 0.0001 |
| LDL-C (mmol/L) | 3.7±0.9 | 3.7±1.0 | 1.000 |
| Family History of T2DM (n %) | 120 (60%) | 55 (18.3%) | 0.0001 |
| Family History of GDM (n %) | 46 (23%) | 13 (4.3%) | 0.0001 |
| R<sub>x</sub> (Diet/Insulin) | 180 (90%)/ 20 (10%) | NA | NA |
| All continuous variables represented by Mean ± standard deviation. Independent sample t-test is done comparing GDM and non-GDM subjects. Categorical variables compared using chi-square analysis. NA= Not applicable/ Not analyzed. |  |  |  |

### Molecular genotype analysis

The genotypes of I28005D polymorphism were in HWE (p>0.05) with the call rates of >95%. The association observed with GDM and non-GDM subjects in ACE variants are tabulated in 2^nd^ table. The wild type genotype frequencies were higher in non-GDM subjects compared to the GDM cases i.e., 57.5% and 45.5% respectively; heterozygous cum homozygous variants were high in GDM cases i.e., 40% and 32% in ID genotypes and 14.5% and 10.5% in DD genotypes. The statistical significance was observed in D allele, ID genotype and Dominant model in GDM cases (D vs I: OR-1.62 [95%CI: 1.09–2.41]; p=0.01; ID vs II: OR-1.58 [95%CI: 1.02–2.42]; p=0.03; DD+ID vs II; OR-1.62 [95%CI: 1.09–2.40]; p=0.01). After adjusting the age and BMI dominant model was associated in GDM cases (DD+ID vs II; OR-1.60 [95%CI: 1.07–2.44]; p=0.04). Combination of GDM with and without obesity subjects were compared with ACE alleles/ genotypes, are tabulated in Table 3. The combination of GDM and Obesity cases were 54.5%, whereas, non-obesity and GDM subjects are 45.5%. The DD genotypes were found to be more in GDM/Obesity cases i.e., 18.3% but none of the allele and genotypes were found to be statistical association [D vs I: OR-1.48 (0.97–2.25); p=0.06 and DD+ID vs II: OR-1.45 (0.83–2.54); p=0.19). However, adjusted sex and BMI also failed to show positive association (DD+ID vs II: OR-1.24 (0.80–2.58); p=0.32).

### Relation between ACE genotypes and glucose values

*ACE* genotypes were compared with weight gain as well as with the biochemical analysis described in detail in table 4. DD genotypes were found to be highly gained the weight, prone to increase BMI when compared with ID and II genotypes (p=0.007 and p=0.02). Comparison between II, ID and DD genotypes showed the relation with glucose values such as FBS, PPBG, GCT and OGTT fasting (p<0.05). The mean of FBS levels were found to be high in ID genotype (4.21±11.25; p=0.0001), PPBG seems to be high in DD genotype (10.64±2.43; p=0.001), GCT was seems to be 2.48±4.22 in DD genotype (p=0.07) and fasting values of OGTT was seems to be high in II genotype as 4.28±2.05 (p=0.0005).

## Discussion

The aim of this current study was to investigate the association between ACE I28005D polymorphism and expansion of GDM in the Saudi population. Results of this study exposed significant association between *ACE* alleles, heterozygous genotype and with dominant model (p<0.05). The inconsistency association occurs in other genotypes with and without adjusting through age and BMI could be attributed small sample size in single exploration and ethnic variations. Genetic etiology with GDM has similar overlap with T2DM; molecular tests for complex diabetes were not beneficial clinically but obliging for MODY disease efficacy for managing family risk and pregnancy treatment.^9^ GDM Incidence was increasing simultaneously with the pandemic in T2DM and obesity in an income developing countries.^8^ The prevalence of GDM in Saudi Arabia was increased from 12–18% and simultaneously, maternal obesity is also increasing in the global world and recent study implemented the stringent guidelines for analysis and diagnosis of GDM incidence increasing burden of Obesity care.^18, 19^ Obesity is known as weight gain or extra fat accumulates and links with the human BMI. Obesity is known as risk factor for T2DM as well as GDM during pregnancy and in this study, relation has been confirmed with obesity and GDM women. Allele and genotype association was not observed when compared between GDM with and without obesity (p>0.05; Table 3). Earlier studies were carried out in different global populations and reported as not associated with D allele with glucose metabolism, T2DM and CAD^20–22^, but when comparison was carried out between weight and BMI with ACE genotypes in GDM subjects, statistical association were observed in Anova analysis (p<0.05; Table 4). Current results are similar with Thailand women.^23^

The relation between T2DM and GDM are still controversy and connection between diabetes and *ACE* have been molded through Renin angiotensin aldosterone system (RAAS), plays a vital role in hypertension and considered as endocrine system and RAAS genes have the correlation between glucose metabolism and hypertension. Angiotensin II appear in RAAS effect glucose metabolism, involved in DM pathogenesis through insulin signal transduction, glucose uptake reduction, β-cells of pancreas inducing oxidative stress. *ACE* is one of genes appear in RAAS group.^24^ There are limited studies performed in GDM with ACE variants; However, studies shown to be both positive and negative association.^12, 25, 26^ Earlier meta-analysis study performed in Arab populations concluded D allele associated with T2DM.^27^ and this current study was in agreement with this meta-analysis report. In Saudi population, limited studies have been performed in various diseases with ACE polymorphism and results concludes both positive/negative association.^13^, ^27–35^ Other meta-analysis reports performed in different diabetic diseases confirm significant association.^36–38^

The strength of this current study was implemented the purely Saudi women to identify the accurate risk in the Saudi population. The non-Saudi women has been excluded as this is a major part of this study.^39^ The another strength of this study were compared the obesity with and without GDM subjects in the Saudi women (Table 3). This case-control study has performed with limited and equal number of sample size i.e., 200 in GDM; 200 in Non-GDM subjects and heterogeneity cannot be excluded as it appears in Saudi population. The limitations of this study could be confirm as (i) skipping of ACE levels (ii) implemented only risk variant (I28005), and (iii) performed limited statistics as this was a case-control study and (iii) skipped to incorporate hypertension values of the pregnant women.

In conclusion, current study confirms the positive association with *ACE* variant in GDM subjects; confirms the relation between GDM and obesity in Saudi women. However, future studies should be implement in other diseases in Saudi as well as GDM and obesity studies in global populations. Meta-analysis studies may give the feedback with *ACE* gene associated with GDM disease.

## Conflict of Interest

None

## Acknowledgement

This study was supported by the college of Medicine Research Centre, Deanship of Scientific Research, King Saud University, Riyadh, Kingdom of Saudi Arabia. I am truly thankful to Prof Alharbi towards his support for this research article.

